# Galectin-3 Mediates Vascular Dysfunction in Obesity by Regulating NADPH Oxidase 1

**DOI:** 10.1101/2023.04.19.537592

**Authors:** Caleb A. Padgett, Róbert K. Bátori, Andrew C. Speese, Cody L. Rosewater, Weston B. Bush, Cassandra C. Derella, Stephen B. Haigh, Hunter G. Sellers, Zachary L. Corley, Madison A. West, James D. Mintz, Brittany B. Ange, Ryan A. Harris, Michael W. Brands, David J. R. Fulton, David W. Stepp

**Affiliations:** Vascular Biology Center, Medical College of Georgia, Augusta University, Augusta, GA; Department of Physiology, Medical College of Georgia, Augusta University, Augusta, GA; Department of Surgery, Medical College of Georgia, Augusta University, Augusta, GA; Department of Pharmacology and Toxicology, Medical College of Georgia, Augusta University, Augusta, GA; Georgia Prevention Institute, Medical College of Georgia, Augusta University, Augusta, GA

**Keywords:** Obesity, Metabolic syndrome, Endothelial Dysfunction, Galectin 3, NADPH Oxidase

## Abstract

**Rationale:** Obesity increases the risk of cardiovascular disease (CVD) through mechanisms that remain incompletely defined. Metabolic dysfunction, especially hyperglycemia, is thought to be a major contributor but how glucose impacts vascular function is unclear. Galectin-3 (GAL3) is a sugar binding lectin upregulated by hyperglycemia but its role as a causative mechanism of CVD remains poorly understood.

**Objective:** To determine the role of GAL3 in regulating microvascular endothelial vasodilation in obesity.

**Methods and Results:** GAL3 was markedly increased in the plasma of overweight and obese patients, as well as in the microvascular endothelium of diabetic patients. To investigate a role for GAL3 in CVD, mice deficient in GAL3 were bred with obese *db/db* mice to generate lean, lean GAL3 knockout (KO), obese, and obese GAL3 KO genotypes. GAL3 KO did not alter body mass, adiposity, glycemia or lipidemia, but normalized elevated markers of reactive oxygen species (TBARS) in plasma. Obese mice exhibited profound endothelial dysfunction and hypertension, both of which were rescued by GAL3 deletion. Isolated microvascular endothelial cells (EC) from obese mice had increased NOX1 expression, which we have previously shown to contribute to increased oxidative stress and endothelial dysfunction, and NOX1 levels were normalized in EC from obese mice lacking GAL3. EC-specific GAL3 knockout mice made obese using a novel AAV-approach recapitulated whole-body knockout studies, confirming that endothelial GAL3 drives obesity-induced NOX1 overexpression and endothelial dysfunction. Improved metabolism through increased muscle mass, enhanced insulin signaling, or metformin treatment, decreased microvascular GAL3 and NOX1. GAL3 increased NOX1 promoter activity and this was dependent on GAL3 oligomerization.

**Conclusions:** Deletion of GAL3 normalizes microvascular endothelial function in obese *db/db* mice, likely through a NOX1-mediated mechanism. Pathological levels of GAL3 and in turn, NOX1, are amenable to improvements in metabolic status, presenting a potential therapeutic target to ameliorate pathological cardiovascular consequences of obesity.

Figure 8.
Visual abstract

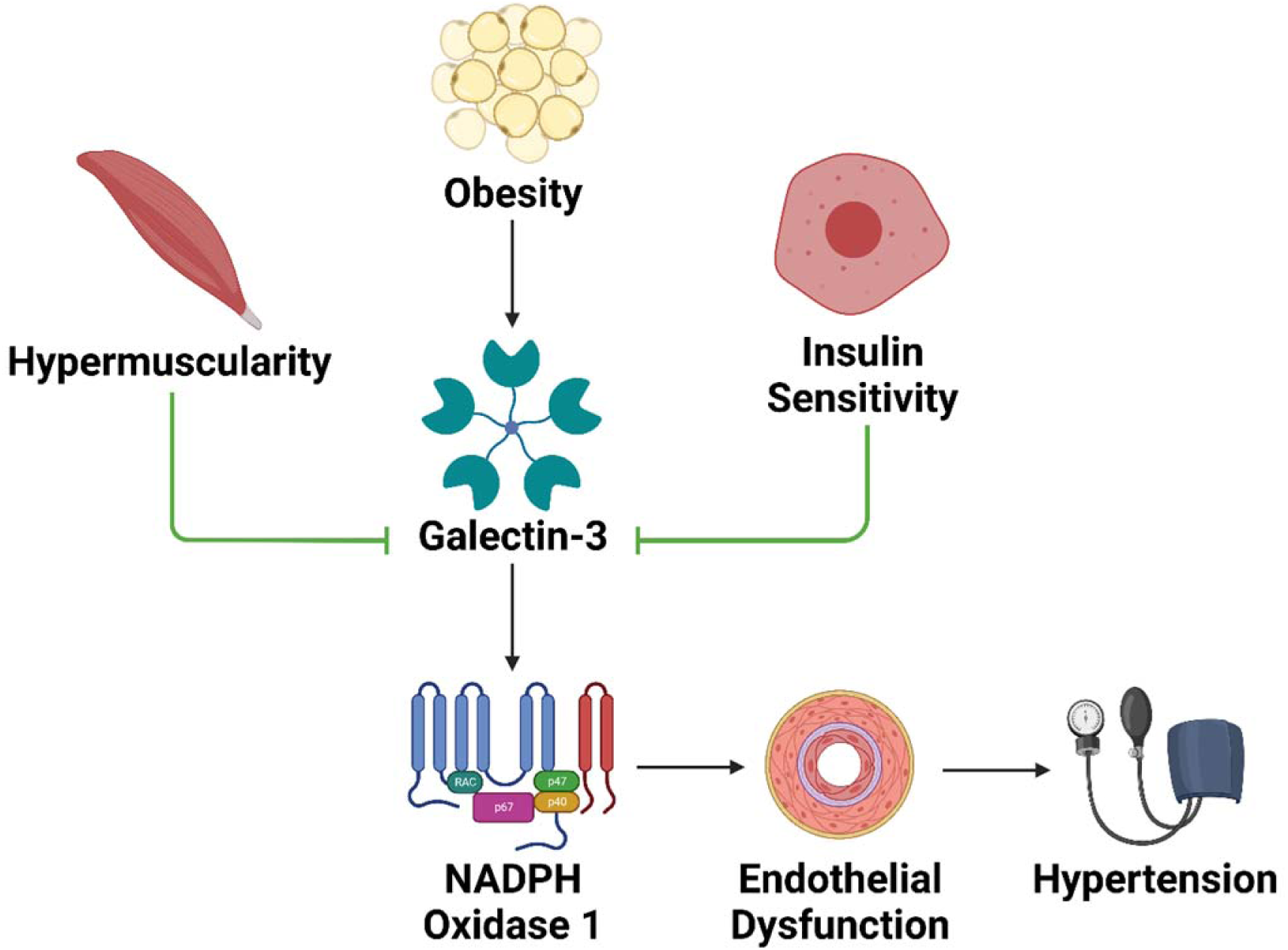

## Introduction

Obesity has long been appreciated as one of the most significant risk factors for cardiovascular diseases, such as hypertension, atherosclerosis, stroke, and myocardial infarction.^1^ Initial dysfunction of the endothelium has been demonstrated to precede these acute cardiovascular events and serves as a mechanistic entry point to later morbidity.^2,3^ Endothelial dysfunction primarily manifests as an inability of the endothelium to effect vasodilation due to a lack of bioavailable nitric oxide (NO).^4^ NO bioavailability is most drastically impaired by reactive oxygen species (ROS), the overproduction of which is a hallmark of vascular disease.^5,6^

Sources of oxidant stress in the vasculature has been a source of intense scrutiny in recent years. We and others have shown that obesity significantly increases ROS levels in the vasculature, with NADPH Oxidase I (NOX1) being the most predominant source of vascular superoxide.^7–11^ Superoxide binds rapidly and inactivates extracellular NO and has further been demonstrated to cause eNOS uncoupling and subsequent production of ROS by eNOS itself, exacerbating lipid peroxidation and protein oxidation.^12–15^ Genetic deletion of NOX1 in obese mice has been shown to be vasoprotective with no improvement in overall metabolism, arguing that NOX is a critical effector linking metabolic dysfunction to impaired vasodilation. However, a significant gap in our knowledge are the upstream mechanisms driving the expression of NOX1 in obesity, which remain incompletely understood and therapeutically elusive.^16,17^

A hallmark of metabolic disease is the inability to properly regulate glucose homeostasis to the detriment of tissues such as the liver, pancreas, and vasculature.^18^ Hyperglycemia has been shown to upregulate the receptor for advanced glycation end-products (RAGE), which, when stimulated by hyperglycated substrates, initiates a swath of proapoptotic and proinflammatory pathways.^19–21^ Many previous studies have examined the role of RAGE in obesity and diabetes, but to date, therapeutics targeting this pathway have proven elusive.^22–24^ Galectin-3 (GAL3), a carbohydrate binding lectin, is a novel form of RAGE studied primarily in the context of cancer, but is highly expressed in hyperglycemia.^25–27^ Interestingly, elevated plasma levels of GAL3 have been proposed as a biomarker for heart disease, as the degree of expression is highly correlated with adverse cardiovascular outcomes.^28–30^ While recognition of GAL3 as a biomarker has increased over several years, the functional role of GAL3 in metabolic and cardiovascular diseases remains incompletely understood, and discovery of a precise mechanism is needed to identify potential therapeutic targets for obese patients with cardiovascular disease.

In the present study, we hypothesized that aberrant glucose regulation in obesity significantly contributes to the upregulation of GAL3 in the endothelium, thereby causing vascular dysfunction and contributing to the development of hypertension and downstream cardiovascular disease. Here we show that GAL3 is a novel regulator of NOX1-derived ROS production in the microvascular endothelium and is an essential metabolic sensor linking metabolic changes in obesity to vascular disease.

## Methods

### Data Availability

Additional data to support this study are available upon reasonable request from the corresponding author.

### Human Studies

Thirty-two healthy men and women reported to the Laboratory of Integrative Vascular and Exercise Physiology (LIVEP) at the Georgia Prevention Institute after completing an overnight fast (no food or drink after midnight). Participants were excluded for any of the following: evidence of cardiovascular disease, renal disease, metabolic disease, rheumatoid arthritis, liver dysfunction, or hypertension. All participants completed the informed consent process. Height and weight were determined using a stadiometer and standard platform scale (CN20, DETECTO, Webb City, MO) and later used for the calculation of body mass index (BMI). Based on BMI, participants were either assigned to a Lean (BMI: 18.5-24.9 kg/m^2^) or Overweight/Obese group (BMI>25.0kg/m^2^). A single stick venous blood draw of the antecubital fossa was obtained at the experimental visit. Blood samples were centrifuged, and plasma separated and stored at −80°C until further analysis. All study protocols were approved by the Institutional Review Board at Augusta University. Pertinent data are included in Supplemental Table 1.

### Animal Studies

All experiments involving animals were conducted under Institutional Animal Care and Use Committee (IACUC) approval and in accordance with the NIH Guide for the Care and Use of Laboratory Animals. Mice were genotyped by isolating DNA from tail clippings at time of weaning with the primers listed in Online Table 1. Animals were sacrificed at approximately 20 weeks of age by being anesthetized in an induction chamber with 5% isoflurane at 1 L/min O_2_ and decapitated by guillotine.

### Animal Model

Mice heterozygous for mutation in the leptin receptor (Jackson Labs; strain no. 000697) were crossed with mice lacking Galectin-3 (Jackson Labs; strain no. 006338) to generate H_db_H_Lgals3_ (lean), H_db_K_Lgals3_ (lean Gal3 KO), K_db_H_Lgals3_ (obese), and K_db_K_Lgals3_ (obese Gal3 KO) genotypes. Endothelial GAL3 KO mice were generated by crossing mice expressing VEC-Cre (Jackson Labs; strain no. 017968) with Lgals3*^fl/fl^*mice (MRC Harwell UK; clone EPD0377_1_A09). 20 week old mice were utilized in each experiment. For metformin studies, 16 week old lean *db^+/-^* and obese *db^−/−^*mice were treated with 5 mg/mL metformin in drinking water for 4 weeks. *db/Nox1, db/Mstn* and *db/Ptp1b* mice were generated as previously described and all mice were genotyped with primers listed in Supplemental Table 2.^7–8,39^

### AAV Production and Administration

Human AgRP (Origene CAT#: RC217144) was cloned into CAG-NLS-GFP (Addgene#: 104061) between KpnI (NEB CAT#: R3142L) and EcoRI (NEB CAT# R3101L) restriction enzyme sites. AAVs were generated following the protocol detailed by Challis *et al.*^31^ Viral titer was determined via qPCR using primers to WPRE (WPRE forward: GGCTGTTGGGCACTGACAAT, WPRE reverse: CCGAAGGGACGTAGCAGAAG). CAG-NLS-GFP AAV.PHP.eB control AAVs were obtained from Addgene (CAT#: 104061-PHP.eB). Obesity or GFP fluorescence was induced via retro-orbital administration of AAV at a dose of 7×10^10gc/mouse under isoflurane anesthesia. For more detailed information regarding the obesity phenotype of this AAV induced model, refer to the associated research letter to this manuscript by Sellers and Padgett *et al*.^32^

### Cell Culture

Microvascular endothelial cells from healthy and Type II diabetic humans (Lonza) were cultured to passage 2-3 in EBM-2 media supplemented with microvascular endothelial cell growth medium SingleQuots supplements (EGM-2 MV) containing 5% fetal bovine serum (FBS). For high glucose and siRNA studies, cells were treated with 15 mM glucose and 10 nM scramble siRNA or GAL3 siRNA for 48 hours. For studies assessing upstream mechanisms, endothelial cells were supplemented with 5% mouse serum for 48 hours. To assess Nox1 promoter activity, cells were transfected with a control plasmid (Bgal), the NOX1 proximal promoter with or without the indicated constructs and a control *Renilla* luciferase.^33–34^ 48 hours later, cells were lysed using a reporter lysis buffer (Promega), and promoter activity was measured by a dual luciferase system using firefly luciferase (Promega) normalized to *Renilla* luciferase (coelenterazine, Nanoliter Technology). Mutant GAL3 constructs were generated as previously described.^35^

### Body Composition

Before euthanasia, mice were weighed and subjected to Nuclear Magnetic Resonance (NMR) in a Minispec Body Composition Analyzer (Bruker; Model no. LF90II) to determine whole-body fat and lean percentages. Heart, lung, spleen, liver, kidney, gonad, and gastrocnemius weights were recorded, as well as snout-anus and tibia lengths. Liver and gastrocnemius tissues were also subjected to NMR to determine intra-organ fat content.

### Metabolic Phenotype

Mice were fasted for 4 hours before euthanasia and blood was collected by heparinized syringe. Blood samples were analyzed for fasting blood glucose using a standard glucometer (AlphaTrak) and for HbA1c using a multi-test A1CNow HbA1c system (PTS Diagnostics). Following whole blood analysis, plasma was separated via centrifugation and analyzed for total cholesterol (FujiFilm), insulin (Alpco), free fatty acids (Sigma), triacylglycerols (FujiFilm), thiobarbituric acid reactive substances (Cayman), and TNFα (BioLegend). Additionally, a cohort of mice were monitored in a Comprehensive Lab Animal Monitoring System (Columbus Instruments) to measure food and water intake, energy expenditure, respiration, and activity. Mice were allowed to acclimate to the CLAMS for 24 hours, then monitored for 72 continuous hours for the purpose of data collection.

### Dihydroethidium & Hydroxyguanosine Staining

Thoracic aortae and mesenteric arteries from euthanized mice were excised, flushed with PBS, and visceral fat trimmed away. Samples were immediately slow-frozen in OCT, cryosectioned, and mounted. Dihydroethidium samples were stained with 2 µM DHE stain (Life Technologies) at 37°C for 30 minutes. Slides were immediately examined using an EVOS FL microscope or a Leica Stellaris confocal microscope. Hydroxyguanosine samples were stained with 1:500 8-OHdG/8 polyclonal antibody (ThermoFisher) overnight, then with 2 µg/mL AlexaFluor 594 secondary antibody (Invitrogen). Samples were imaged using a Leica Stellaris confocal microscope.

### Endothelial Cell Isolation

Thoracic aortae and mesenteric arteries from euthanized mice were excised, flushed with PBS, and visceral fat trimmed away. Arteries were minced and incubated in dispase/collagenase II digestion buffer at 37°C for 1 hour. Homogenates were centrifuged, cells resuspended in endothelial cell isolation buffer, and incubated with CD31 microbeads (Miltenyi Biotec; Lot no. 5191031675) at 4°C for 20 minutes. Cells were then washed with endothelial cell isolation buffer and applied to a magnetic column where flow through fraction was collected. Following wash of column with 2 mL buffer, column was flushed with 1 mL endothelial isolation buffer to collect endothelial fraction. Cells were spun down and resuspended in Trizol for RNA isolation or Laemmli buffer for western blot. Endothelial cell fractions were assessed for purity by qPCR (Supplemental Figure 1).

### Gene Expression

RNA was isolated from endothelial and flow through fractions using Direct-Zol RNA MiniPrep Plus (Zymo; Lot no. ZRC204808). cDNA was synthesized using OneScript cDNA Synthesis SuperMix (ABM; cat no. G452). qPCR was performed in a CFX-Connect Real-Time PCR Detection System (Bio-Rad) utilizing BrightGreen Express 2x qPCR MasterMix - iCycler (ABM; cat no. MasterMix-EC). Products were amplified at 95°C for 3 minutes followed by cycles of 95°C for 15s, 58.5°C for 15s, and 72°C for 15s for 40 cycles. Expression fold change was calculated using the 2^−ΔΔCT^ method normalized to 18s rRNA as an internal control. Primer sequences are found in Supplemental Tables 3-4.

### Protein Expression

Isolated endothelial cells were suspended in Laemmli (1X) buffer and boiled at 95°C for 5 minutes. Samples were run on a 10% SDS-Page gel for 2 hours at 100V. Separated proteins were then transferred to an PVDF membrane activated by soaking in methanol for 30s, then placed in transfer buffer (100 ml Transfer Solution (144 g Glycine, 30.2 g Tris-Base to 1,000 L), 200 ml 100% methanol, 700 ml DI H_2_O) for 2 minutes prior to transfer. Transfer was performed at 4C for 1 hour and 15 minutes at 100V. Membranes were blocked in 5% milk for 1 hour and washed for 15 minutes 3 times with 1X TBST. The membrane was then cut to separate the desired proteins for probing. Membrane slices were probed with their respective antibody (Supplemental Tables 4-5) overnight at a dilution of 1:1000 in 5% Milk, unless otherwise specified by the manufacturer. The membranes were then washed again as above. Membranes were probed with secondary antibody at a dilution of 1:10,000 in 5% milk and incubated at 25°C for a maximum of 1 hour, then washed again as previously described. ECL was used to activate the secondary antibodies, per manufacturer’s instructions. The strips were lined up accordingly as to appear prior to cutting, then gently dabbed with Kim-wipes to remove any excess ECL. Activated membranes were then imaged via chemiluminescence with a 10-30 second exposure. Samples were analyzed by densitometry and normalized to loading control, then reran with normalized values. Normalized samples were then prepared and imaged as above.

### Wire Myography

Thoracic aortae were dissected, fat trimmed, and sliced into rings for mounting on a wire myograph (DMT) in Krebs solution at 37°C. Vessels were assessed for viability with saturated KCl solution, washed in Krebs solution and subjected to dose response curves (10^−9^ – 10^−4^ M) of acetylcholine, phenylephrine, and sodium nitroprusside, with washes in between each curve.

### Pressure Myography

Second and third order mesenteric arteries were dissected from intestines in ice-cold Krebs solution (118 mM NaCl, 25 mM NaHCO_3_, 11.1 mM d-glucose, 4.71 mM KCl, 2.56 mM CaCl_2_, 1.13 mM NaH_2_PO_4_, 7 mMMgCl_2_). Dissected arteries were trimmed free from adipose tissue, cannulated, and secured on glass pipettes in 10 mL of Krebs solution in a single vessel chamber (Living Systems Instrumentation). Microvessels were pressurized to physiological pressure of 60 mmHg with a Pressure Servo Controller (LSI; Model no. PS-200-S) and heated to physiological temperature of 37°C with a Temperature Controller (LSI; Model no. TC-095). Vessels were allowed to equilibrate for 20 minutes, after which viability was assessed with 10 µL of saturated KCl. After 3 washes to ensure KCl removal, dose-response curves of acetylcholine, phenylephrine, and sodium nitroprusside (10^−9^ – 10^−4^ M) were generated, with 3 washes in between each of the curves. Once dose-response curves were generated, vessel was washed 3 times in calcium-free Krebs solution and allowed to equilibrate for 60 minutes. Myogenic tone was assessed at 20 mmHg increments (20 – 120 mmHg) by recording passive inner and outer diameters at each pressure.

### Blood Pressure Telemetry

Telemetry experiments were performed as previously described.^40^ Briefly, mice were anesthetized and implanted with DSI telemetry units via the left carotid artery. Mice were allowed to recover for approximately 7 days, then blood pressure was recorded 24 hours per day for 3 days.

### Statistical Analyses

Analysis of association between BMI and GAL3 was performed with a simple linear correlation. A student’s T-test was calculated to examine differences between two groups. For experiments with lower n, the exact versions of non-parametric tests were used as they are appropriate tests to use with smaller sample sizes. Mann-Whitney tests were calculated to examine differences between two groups, and exact Kruskal-Wallis tests were calculated to examine differences among four groups. All analyses were performed with GraphPad Prism 9.1 (GraphPad Software Inc.). Data are expressed as mean ± standard error of the mean. P≤0.05 was used as criteria for significance. **** = p<0.0001, *** = p<0.001, ** = p<0.01, * = p<0.05, ns = not significant.

## Results

### GAL3 is increased in the plasma and microvascular endothelium during diabetes and obesity

Plasma levels of GAL3 were assessed in lean and obese humans and found to be both significantly upregulated in overweight and obese individuals (Figure 1A) and associated with increasing BMI (Figure 1B). Additionally, GAL3 expression in human microvascular endothelial cells was assessed at both the mRNA (Figure 1C) and protein (Figure 1D) levels and found to be significantly upregulated in the microvascular endothelium from patients with Type II Diabetes Mellitus. Conduit (aorta) and resistance (mesenteric) arteries were isolated from 20-week old lean (*db/+*) and obese (*db/db*) mice. Endothelial cells from each of these arteries were isolated, assessed for purity (Supplemental Figure 1), and assessed for GAL3 expression. GAL3 mRNA expression was moderately increased in the aortic endothelium of obese male mice (Figure 1E). However, in the microvascular endothelium, GAL3 was increased approximately 160-fold in obese male mice compared to lean controls. Further analysis of the non-endothelial fraction, which includes vascular smooth muscle, fibroblasts, and adipocytes, revealed no significant changes in GAL3 expression, suggesting that major increases in expression result from changes in the endothelium (Supplemental Figure 2A). These results were further verified by Western blot (Figure 1F), indicating that GAL3 is most highly expressed in the microvascular endothelium in obesity. Since the microvasculature is essential for the maintenance of blood pressure and tissue perfusion, we chose to focus on the role of GAL3 in the microvascular endothelium.^36–37^ Assessment of endothelial GAL3 expression in lean and obese female mice revealed no significant upregulation (Supplemental Figure 2B), therefore male mice were utilized exclusively from this point forward in the study.

**Figure 1.**
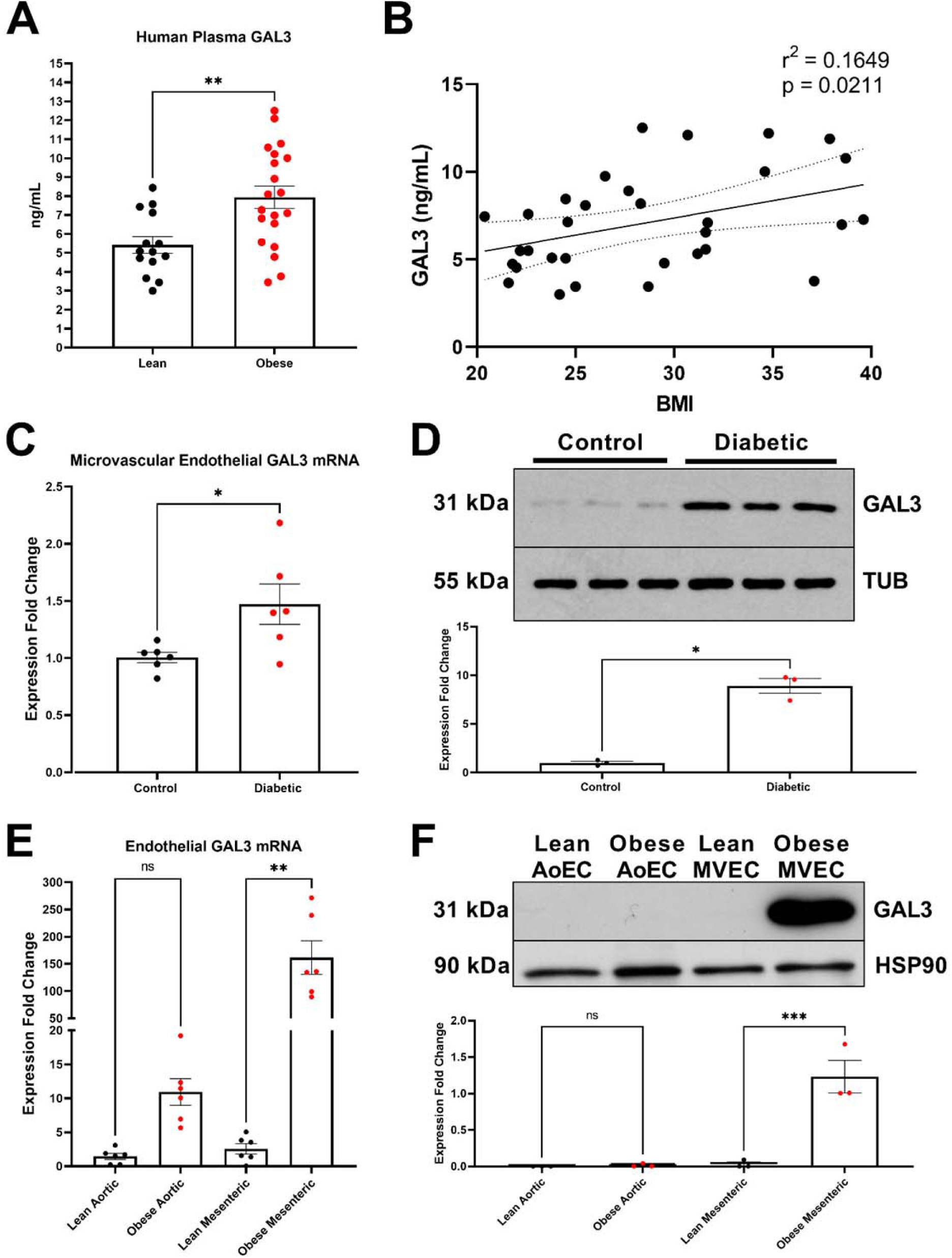
GAL3 expression is increased in the microvascular endothelium in obesity. **A.** Plasma GAL3 levels from lean and obese patients (n=14-20). **B.** Human plasma GAL3 levels according to BMI (n=34). **C.** GAL3 mRNA levels in microvascular endothelial cells from control and diabetic patients (n=6). **D.** Representative Western blot of GAL3 protein levels in microvascular endothelial cells from control and diabetic patients (n=3). **E.** GAL3 mRNA levels in freshly isolated aortic and mesenteric flow through fractions from male 20-week old lean and obese mice (n=6). **F.** Representative Western blot of GAL3 protein levels in freshly isolated aortic and mesenteric endothelial cells from male 20-week old lean and obese mice. All data are represented as mean ± SEM.

### Deletion of GAL3 has no impact on the anatomic or metabolic consequences of obesity

In order to elucidate the mechanistic role of GAL3 in obesity-induced vascular disease, we crossed *db/+* mice with *Lgals3* knockout mice to generate four genotypes: lean (H*_db_*H*_Lgals3_*), lean GAL3 KO (H*_db_*K*_Lgals3_*), obese (K*_db_*H*_Lgals3_*), and obese GAL3 KO (K*_db_*K*_Lgals3_*) (Figure 2A). Heterozygous mice were used as controls in order to reduce the number of animals needed for the study, since *db/db* mice are infertile and must be bred as heterozygotes.^38^ Obese mice exhibited higher body weight (Figure 2B) and fat mass (Figure 2C) compared to lean mice, with GAL3 deletion having no effect on these measures in lean or obese mice. Other indices of body size such as snout-anus length and tibia length were similar in all four genotypes (Supplemental Figure 3). Organ weights were consistent with previous findings in *db/db* mice, including hepatomegaly, renal hypertrophy and muscle atrophy (Supplemental Figure 4).^39^ All other organ weights remained unchanged, and deletion of GAL3 caused no detectable changes between groups (H*_db_*H*_Lgals3_* to H*_db_*K*_Lgals3_* and K*_db_*H*_Lgals3_* to K*_db_*K*_Lgals3_*). All four genotypes were subjected to analysis in CLAMS for further metabolic analysis (Supplemental Figure 5). Obese mice exhibited decreased VO_2_ and VCO_2_ due to inactivity, but did not exhibit differences in respiratory exchange ratio, or heat production. Obese mice also exhibit markedly increased foot and water intake, and markedly decreased wheel activity. Deletion of GAL3 had no effect on any of the parameters assessed by CLAMS (Supplemental Figure 5). Plasma analyses indicated significant increases in HbA1c (Figure 2D), insulin (Figure 2E), total cholesterol (Figure 2F), and free fatty acids (Figure 2G) in obese mice, with GAL3 deletion again causing no significant changes between groups.

**Figure 2.**
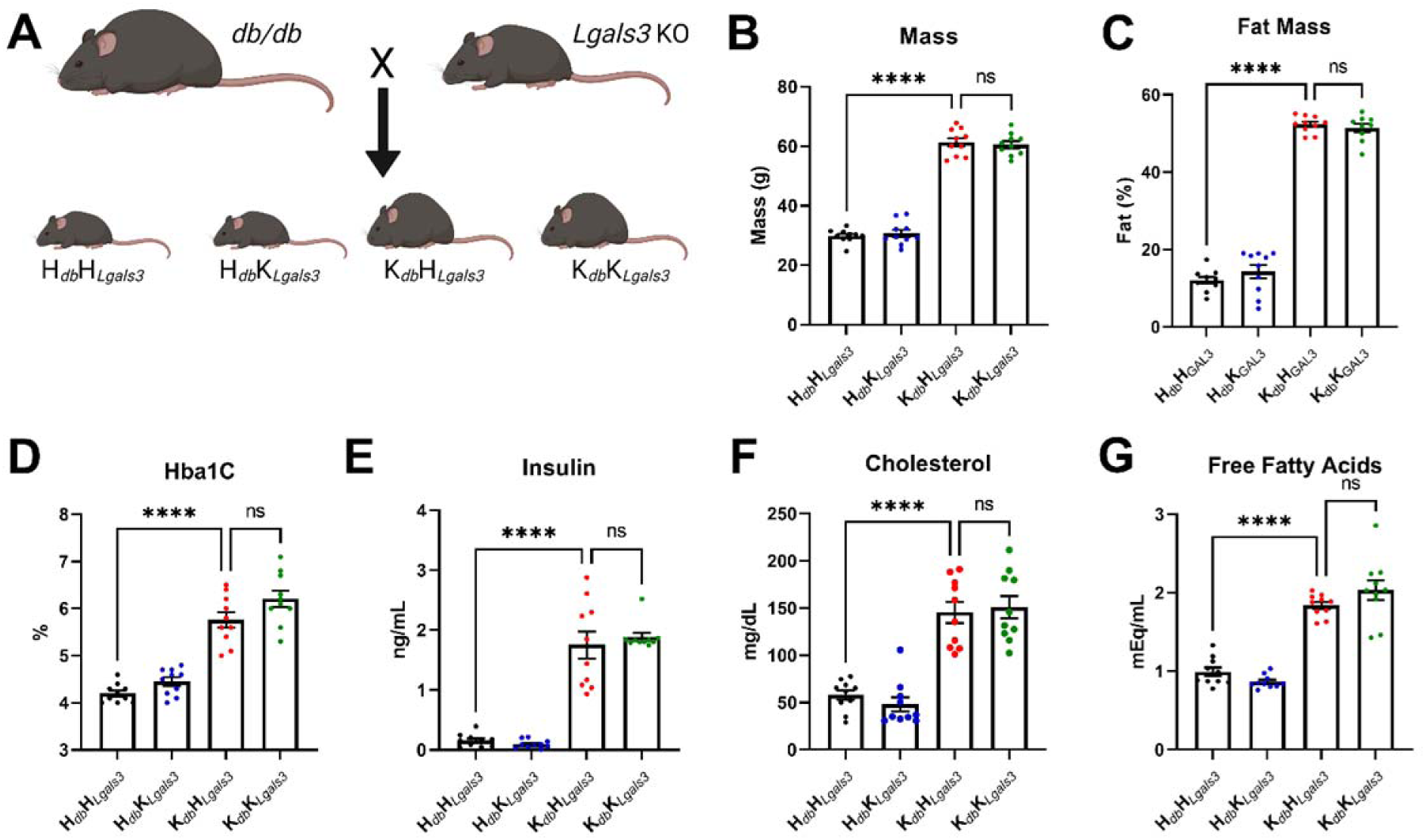
Deletion of GAL3 does not affect glycemia or lipidemia but improves plasma reactive oxygen species production in obesity. **A.** Visual representation of breeding schema. **B.** Total body mass from HH, HK, KH, and KK mice (n=10). **C.** Fat mass from all four genotypes (n=10). **D.** HbA1c from all four genotypes (n=10). **E.** Plasma insulin from all four genotypes (n=10). **F.** Plasma cholesterol from all four genotypes (n=10). **G.** Plasma non-esterified fatty acids from all four genotypes (n=10).

### Deletion of GAL3 improves microvascular endothelial function and hypertension

We next tested the role of GAL3 in contributing to obesity-induced endothelial dysfunction. Aortae and second-third order mesenteric arteries from all four genotypes of mice were subjected to wire and pressure myography, respectively. After preconstriction with phenylephrine, vessels from obese mice exhibited marked endothelial dysfunction as evidenced by a decrease in the vasodilatory response to acetylcholine (Figure 3A,D). However, deletion of GAL3 in obese mice rescued endothelium-dependent vasodilation, as K*_db_*K*_Lgals3_* mice had no significant differences in vasodilatory response to acetylcholine compared to H*_db_*H*_Lgals3_*and H*_db_*K*_Lgals3_* controls (Figure 3A,D). Obese mice exhibited no difference in alpha-adrenergic-dependent vasoconstriction in response to a dose-response curve of phenylephrine (Figure 3B,E), and no differences in endothelial-independent dilation in response to increasing doses of the direct NO donor, sodium nitroprusside (Figure 3C,F). Deletion of GAL3 had no effect on alpha-adrenergic constriction or endothelial-independent dilation in either lean or obese mice. Maximum constriction to potassium chloride, vascular tone, and resting internal diameter were not different among the four groups of mice (Supplemental Figure 6). In order to further investigate the role of GAL3 in cardiovascular disease, we implanted mice with radiotelemetry probes to assess mean arterial pressure (MAP) and heart rate. Obese mice displayed a significant increase in MAP, which was ameliorated by deletion of GAL3 (Figure 3G) and independent of changes in heart rate (Figure 3J). GAL3 deletion did not significantly decrease systolic blood pressure (Figure 3H) but did significantly decrease diastolic blood pressure in obese mice (Figure 3I). Taken together, these data confirm that obesity causes endothelial dysfunction and hypertension, and that GAL3 deletion rescues vasodilatory capacity independent of smooth muscle- or non-endothelial-mediated mechanisms and improves obesity-associated hypertension.

**Figure 3.**
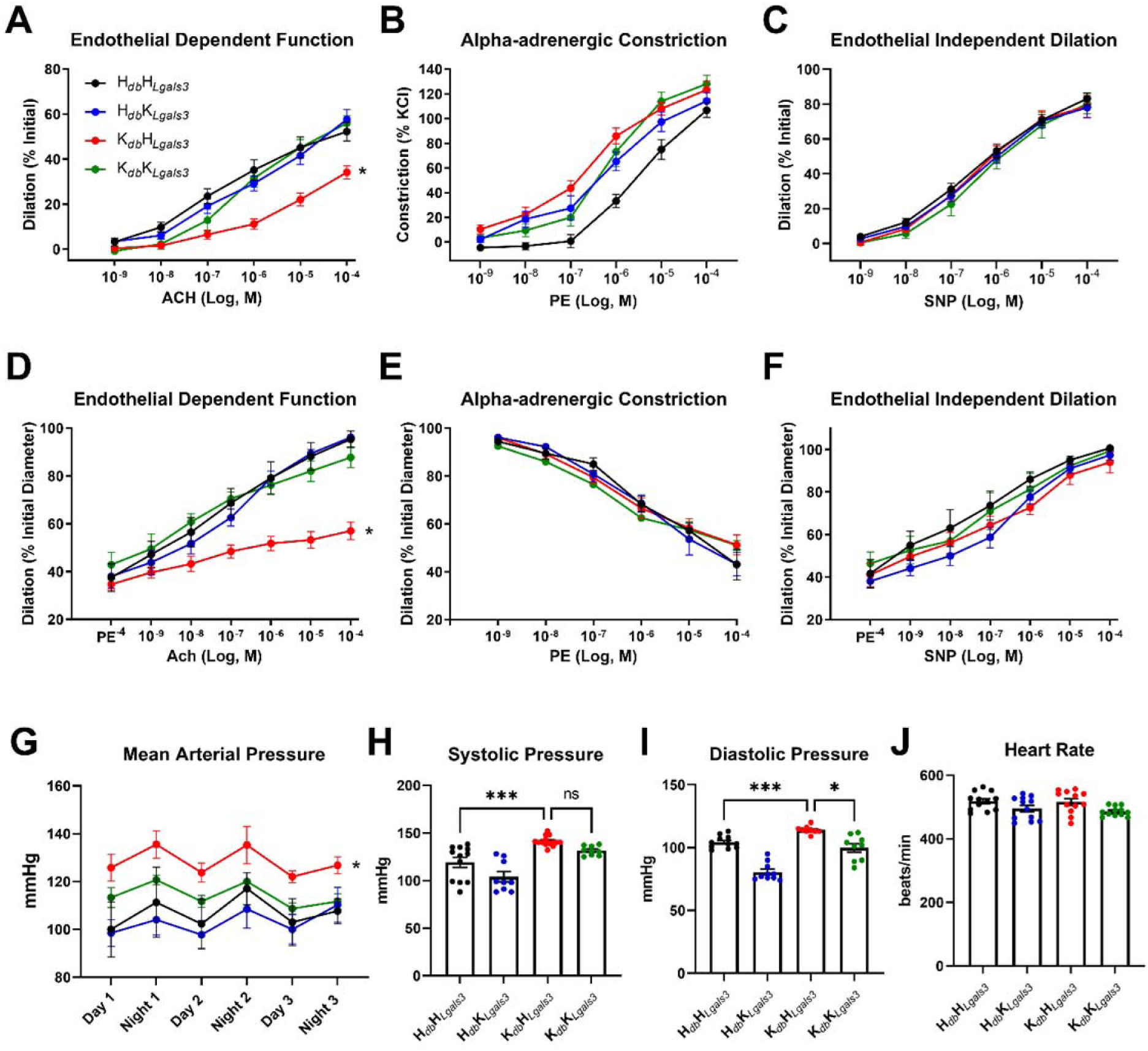
Deletion of GAL3 improves endothelial-dependent dilation in obese mice. **A.** Aortic vasodilation (calculated as percent of initial preconstriction) to dose-response curve of acetylcholine in HH, HK, KH, and KK mice (n=4-6). **B.** Aortic alpha-adrenergic vasoconstriction (calculated as percent of max constriction to KCl) to dose-response curve of phenylephrine in all four genotypes (n=4-6). **C.** Aortic vasodilation (calculated as percent of initial preconstriction) to dose-response curve of sodium nitroprusside in all four genotypes (n=4-6). **D.** Mesenteric arterial vasodilation (calculated as percent of initial diameter) to dose-response curve of acetylcholine in HH, HK, KH, and KK mice (n=4-6). **E.** Mesenteric arterial alpha-adrenergic vasoconstriction (calculated as percent of internal diameter) to dose-response curve of phenylephrine in all four genotypes (n=4-6). **F.** Mesenteric arterial vasodilation (calculated as percent of initial diameter) to dose-response curve of sodium nitroprusside in all four genotypes (n=4-6). **G.** Average mean arterial pressure over 72 hours in all four genotypes (n=3-4). **H.** Average systolic blood pressure over 72 hours in all four genotypes (n=3-4). **I.** Average diastolic blood pressure over 72 hours in all four genotypes (n=3-4). **J.** Average heart rate over 72 hours in all four genotypes (n=3-4).

### GAL3 regulates NOX1-mediated ROS production in the vasculature

Strikingly, deletion of GAL3 normalized plasma thiobarbituric acid reactive substances (TBARS) in obese mice, indicating that GAL3 is a potent regulator of oxidative stress (Figure 4A). Aortic segments from obese mice exhibited higher oxidant stress as evidenced by increased DHE staining, while obese mice lacking GAL3 displayed decreased staining, demonstrating that GAL3 regulates functional ROS production in the vasculature (Figure 4B). Additionally, mesenteric arteries from obese mice demonstrated ROS accumulation as evidenced by increased DHE and 8-OHdG staining, while obese mice lacking GAL3 displayed decreased staining, further demonstrating that GAL3 regulates vascular ROS production in the microvasculature (Figure 4C-D). To investigate the mechanism by which GAL3 deletion improves vascular function, we isolated microvascular endothelial cells from H*_db_*H*_Lgals3_*, H*_db_*K*_Lgals3_*, K*_db_*H*_Lgals3_,* and K*_db_*K*_Lgals3_* mice. Endothelial GAL3 expression was markedly increased in the microvasculature of obese mice, consistent with that shown previously (Figure 4E). NOX1, the major ROS-generating enzyme in the vasculature, was found to be highly upregulated in obesity (Figure 4F), as we and others have previously shown.^7,8^ However, deletion of GAL3 in obese mice normalized increased levels of NOX1 mRNA (Figure 4F) and protein expression (Figure 4G) to that of lean controls, demonstrating that GAL3 is a novel regulator of NOX1 expression. The levels of other vascular NOX isoforms (NOX2 and NOX4) were unaltered by GAL3 deletion, suggesting that GAL3 specifically regulates NOX1 in the endothelium (Supplemental Figure 7). Furthermore, no compensatory upregulation of other RAGEs was observed with deletion of GAL3 (Supplemental Figure 8).

**Figure 4.**
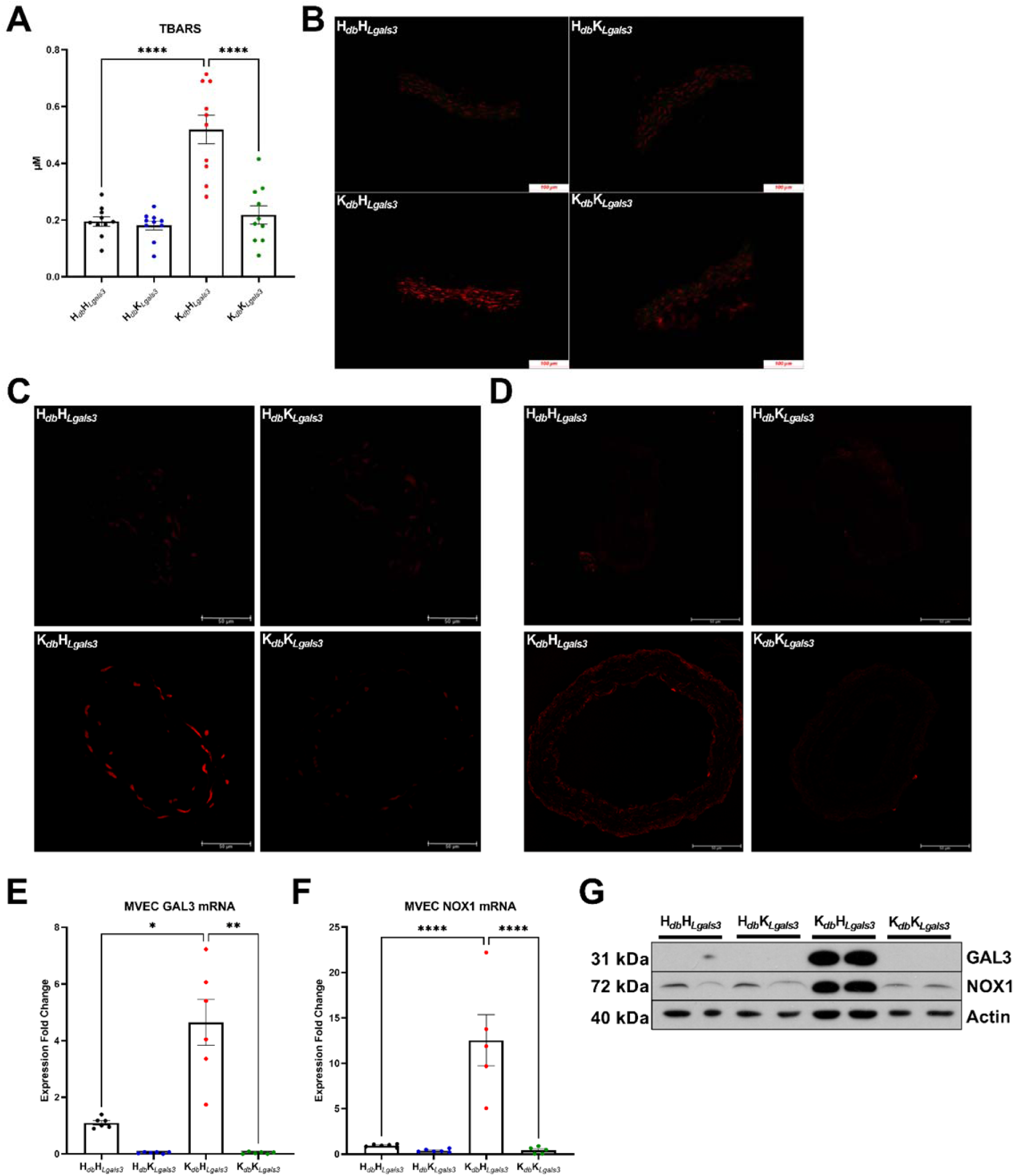
GAL3 regulates endothelial NOX1 expression and reactive oxygen species production. **A.** Plasma TBARS levels from male 20-week old HH, HK, KH, and KK mice (n=10). **B.** Representative DHE staining in aortic segments from all four genotypes. **C.** Representative DHE staining in mesenteric arteries from all four genotypes. **D.** Representative 80OHdG staining in mesenteric arteries from all four genotypes. **E.** GAL3 mRNA levels in freshly isolated mesenteric endothelial cells from all four genotypes (n=5). **F.** NOX1 mRNA levels in freshly isolated mesenteric endothelial cells from all four genotypes (n=5) **G.** Representative western blot of GAL3, NOX1, and HSP90 protein levels in freshly isolated mesenteric endothelial cells from all four genotypes (n=2). All data are represented as mean ± SEM. Scale bar = 100 µm.

Because GAL3 has previously been shown to be pathologically overexpressed in multiple vascular cell types, we generated endothelial cell-specific knockout mice in order to critically test the hypothesis that endothelial cell GAL3 is a key mediator of microvascular dysfunction in obesity.^45^ *Lgals3^fl/fl^*mice were crossed with VE-Cadherin-CRE mice and administered a novel capsid engineered AAV expressing the oreixigen, agouti-related peptide (AgRP) in the brain. Mice administered AgRP AAV subsequently developed hyperphagia with resulting obesity over a 12-week period (Figure 5A).^32^ Consistent with findings in obese db/db mice, GAL3 was significantly upregulated in the microvascular endothelium of AgRP-AAV obese animals and was ablated in obese animals expressing Cre recombinase restricted to the endothelium (Figure 5B). Concurrently, the specific loss of endothelial GAL3 led to a corresponding decrease in NOX1 overexpression, confirming previous results in whole-body GAL3 knockout mice, and advancing a new concept that in the microvasculature, endothelial GAL3 mediates NOX1 overexpression in obesity (Figure 5C). The endothelial specific deletion of GAL3 also rescued obesity-induced microvascular endothelial dysfunction in AgRP-AAV mice (Figure 5D). AgRP-AAV administration and endothelial GAL3 deletion had no appreciable effect on smooth muscle contractility (Figure 5E) or endothelium-independent vasodilation (Figure 5F), with no alterations in maximum constriction, vascular tone, or resting internal diameter (Supplemental Figure 9). These data confirm that GAL3 is a novel regulator of NOX1-derived oxidative stress in the microvascular endothelium, and that targeting GAL3 has the therapeutic potential to abrogate early endothelial dysfunction in obesity.

**Figure 5.**
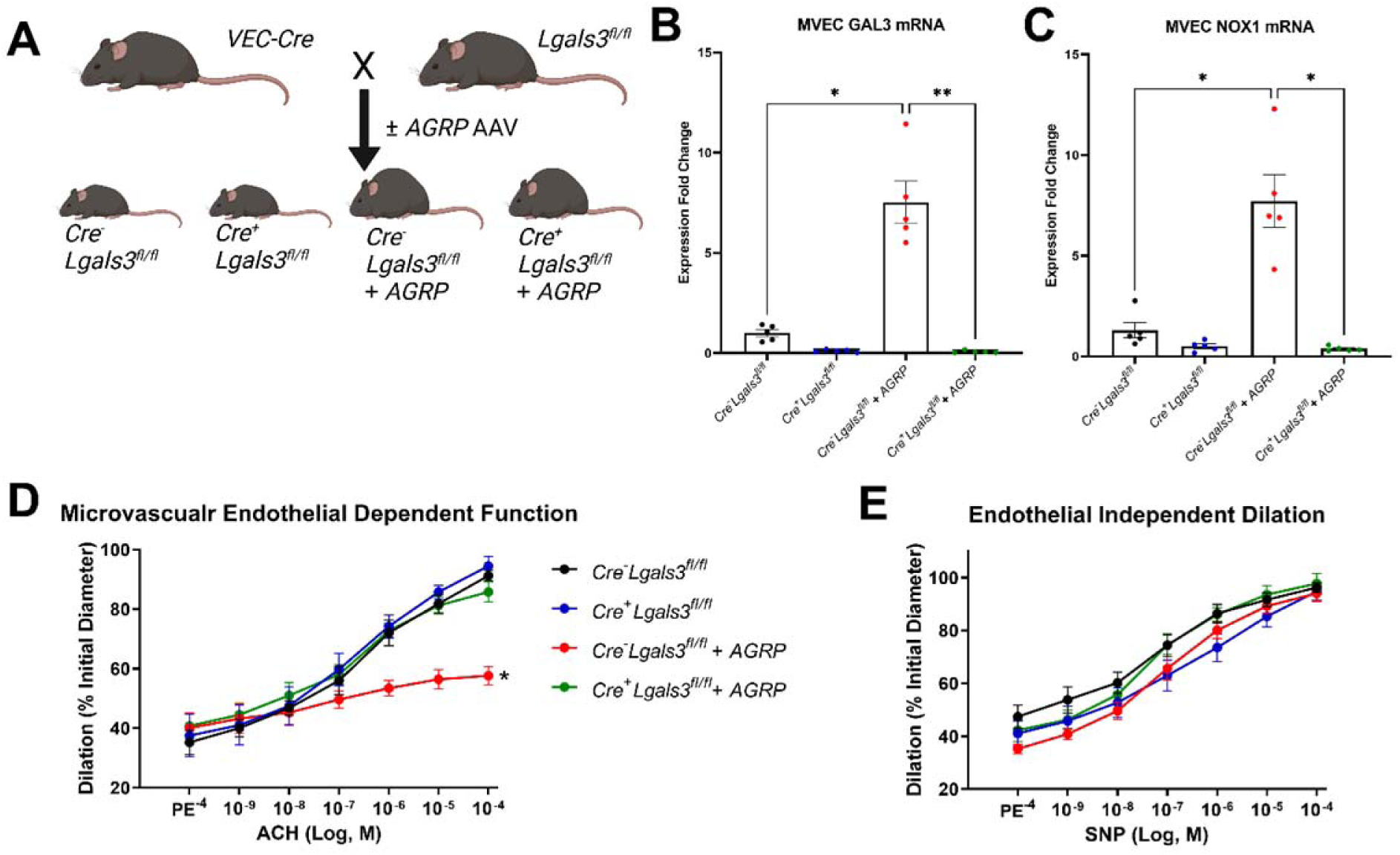
Endothelial GAL3 regulates NOX1-associated endothelial dysfunction in obese mice. **A.** Visual representation of breeding schema. **B.** GAL3 mRNA levels in freshly isolated mesenteric endothelial cells from all four genotypes (n=5). **C.** NOX1 mRNA levels in freshly isolated mesenteric endothelial cells from all four genotypes (n=5). **D.** Mesenteric arterial vasodilation (calculated as percent of initial diameter) to dose-response curve of acetylcholine in all four genotypes (n=4-5). **E.** Mesenteric arterial vasodilation (calculated as percent of initial diameter) to dose-response curve of sodium nitroprusside in all four genotypes (n=4-5). All data are represented as mean ± SEM.

### Altered glycemic metabolism and oligomerization of GAL3 is required for NOX1 expression

We next evaluated the translational significance of our findings in human microvascular endothelial cells *in vitro*. Human endothelial cells were treated with serum isolated from lean or obese mice to test the hypothesis that endothelial GAL3/NOX1 expression is responsive to the hyperglycemic milieu occurring in obesity. Serum from lean mice evoked no expression changes in GAL3 or NOX1, while serum from obese mice significantly increased both GAL3 and NOX1 levels after only 48 hours of treatment (Figure 6A). Treatment with only high glucose also evoked GAL3 and NOX1 overexpression, confirming that the GAL3/NOX1 axis is upregulated by hyperglycemia (Figure 6B). In order to confirm that endothelial GAL3 regulates NOX1 *in vitro* as *in vivo*, GAL3 expression was silenced using siRNA. Human microvascular endothelial cells transfected with GAL3 siRNA failed to upregulate NOX1 in response to high glucose, indicating that GAL3 contributes to NOX1 overexpression in the endothelium by acting as a sensor of hyperglycemia. In order to further probe the interaction between GAL3 and NOX1, we assessed the activity of the NOX1 proximal promotor in GAL3-knockout HEK cells harboring a luciferase reporter. GAL3 transfection significantly increased NOX1 promotor activity, indicating that GAL3 can regulate NOX1 expression at the transcriptional level (Figure 6C). GAL3 is a chimeric galectin that functions through the formation of oligomers. To test the hypothesis that oligomerization is required for GAL3-mediated NOX1 promotor activation, we generated a L131A/L203A double mutant of GAL3 that inhibits its ability to oligomerize. Transfection of GAL3 KO HEK293 cells with wild type GAL3 evoked NOX1 promotor activity similar to previous experiments, but transfection of an inactive mutant of GAL3, that cannot oligomerize, resulted in no difference in NOX1 promotor activity compared to the transfection control. These data indicate that GAL3 regulates NOX1 expression in hyperglycemia and that GAL3 oligomerization is required for NOX1 promotor activity.

**Figure 6.**
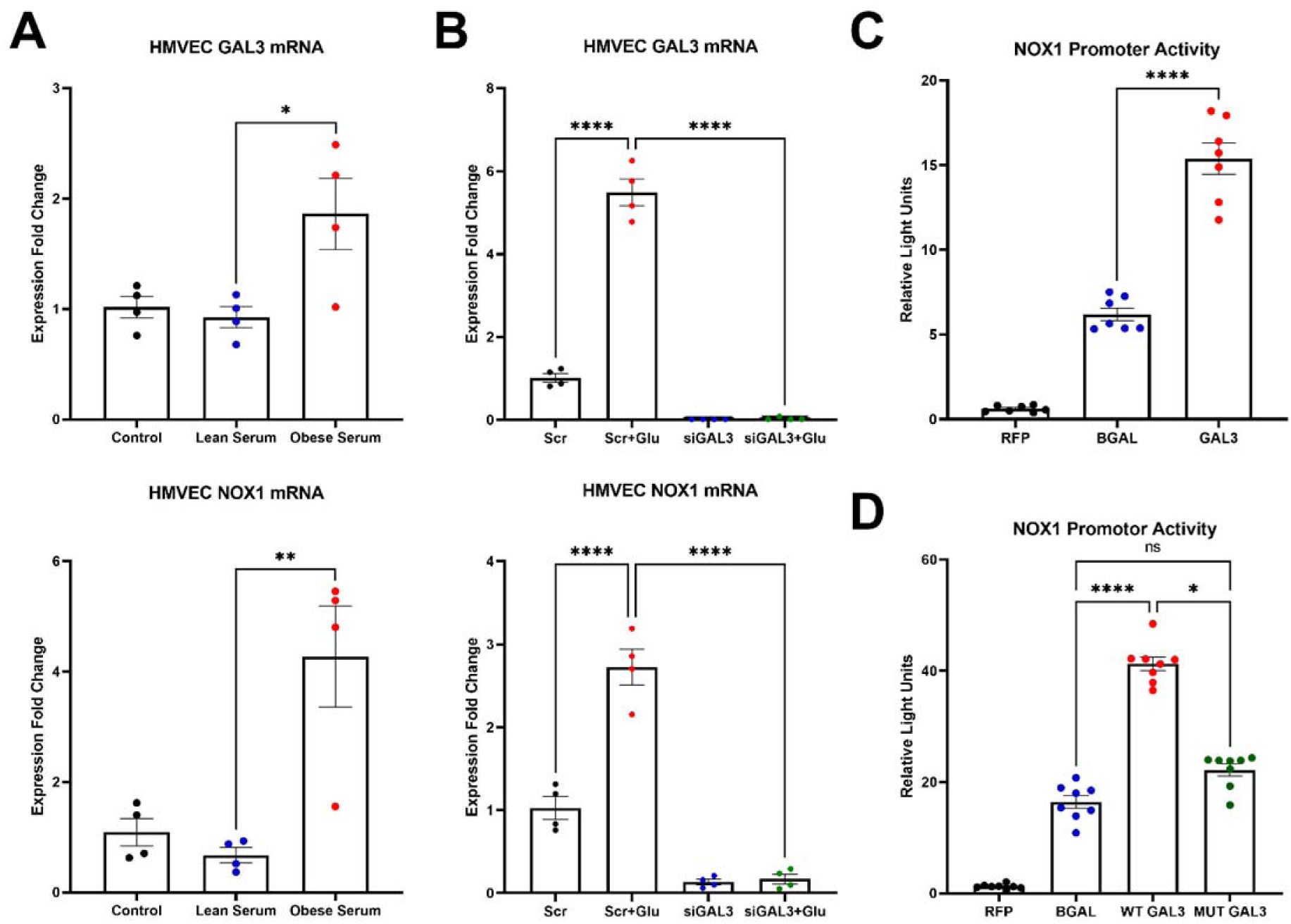
GAL3 regulates NOX1 expression in hyperglycemia. **A.** GAL3 and NOX1mRNA levels in human microvascular endothelial cells treated with lean (mean BG=125.9±10.98) or obese (mean BG=257.1±19.72) mouse serum (n=4). **B.** GAL3 and NOX1mRNA levels in human microvascular endothelial cells treated with high glucose with or without GAL3 siRNA (n=4). **C.** NOX1 promotor activity in GAL3 KO HEK cells transfected with RFP, BGAL, or GAL3 (n=7). **D.** NOX1 promotor activity in GAL3 KO HEK cells transfected with RFP, BGAL, wild type GAL3, or mutated GAL3 (n=8). All data are represented as mean ± SEM.

### Metabolic status is the primary determinant of GAL3 and NOX1 expression

Since amelioration of GAL3 in the endothelium proved beneficial in maintaining a healthy vasodilatory response in obesity, we explored upstream mechanisms of increased levels of endothelial GAL3. Using our previously described mouse model of NOX1 deletion in obesity^7^, we confirmed that GAL3 levels in the endothelium of obese mice are not affected by NOX1 deletion (Figure 7A), indicating that GAL3 is upstream of NOX1 in vivo, and is therefore a suitable target for decreasing NOX1-mediated ROS production in disease by altering the metabolic sensor that regulates NOX1 expression. We next utilized our previously described mouse model of hypermuscularity^8,40–41^ to demonstrate that preservation of muscle mass and glucose metabolism in obesity decreases endothelial GAL3 and subsequent NOX1 levels (Figure 7B). Previous studies from our laboratory have also established that deletion of protein tyrosine phosphatase 1b (PTP1B), a potent negative regulator of insulin signaling, confers protection from obesity-induced endothelial dysfunction.^39^ In the microvascular endothelium, GAL3 and NOX1 levels were normalized to that of lean controls in obese mice lacking PTP1B (Figure 7C). Turning next to a pharmacological approach, we treated lean and obese mice with metformin for 4 weeks, and observed a corresponding decrease in endothelial GAL3 and NOX1 levels (Figure 7D). Changes in GAL3 and NOX1 expression in metformin-treated mice correlated with significantly decreased HbA1c in obese mice with no corresponding decrease in body weight. Taken together, these data indicate that elevated endothelial GAL3 expression in obesity can be reduced to normal levels by improvements in metabolism, either through the preservation and augmentation of skeletal muscle mass, improvement of insulin signaling, or enhanced glucose handling through treatment with metformin.

**Figure 7.**
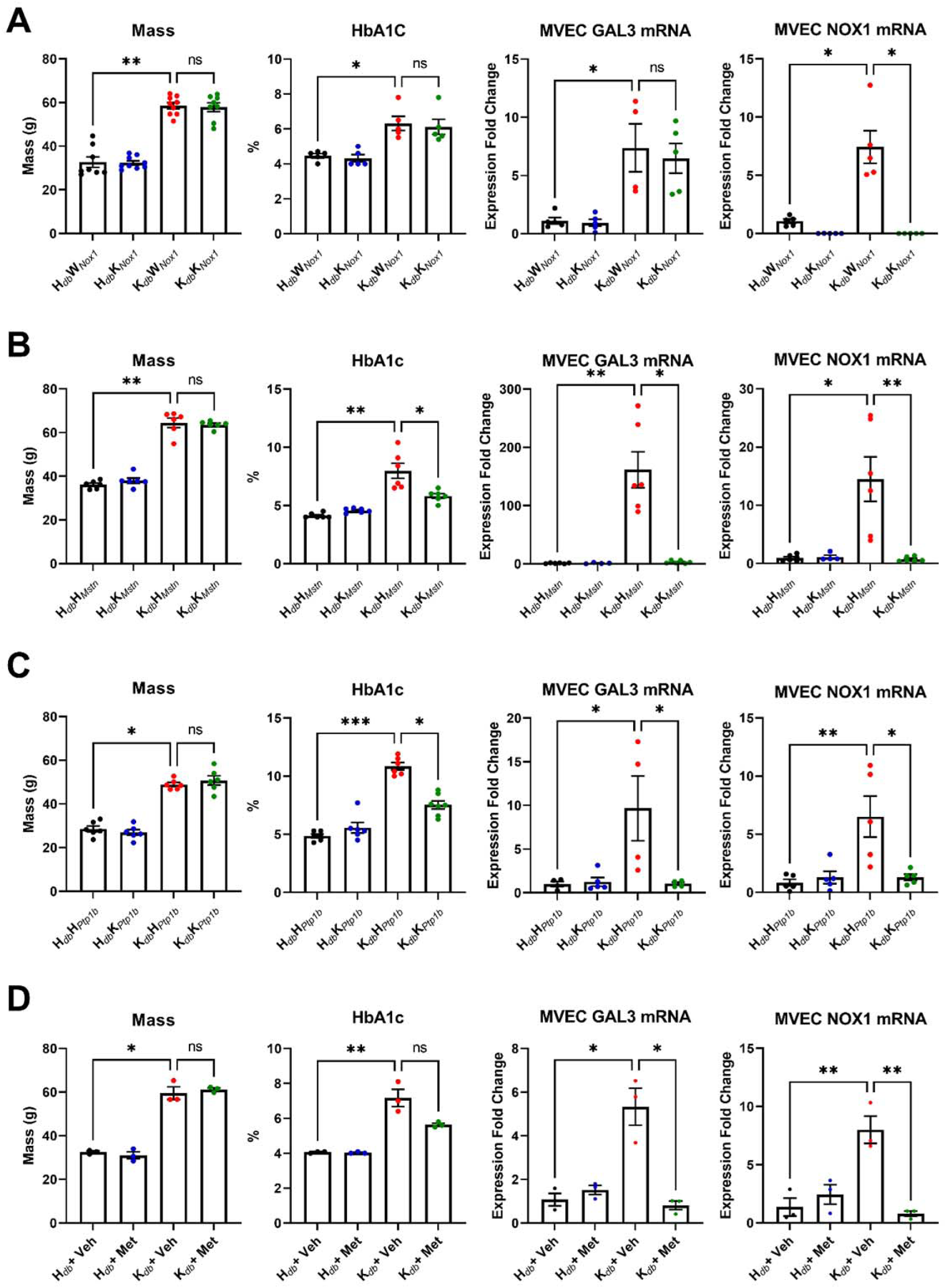
GAL3 is upstream of NOX1 and is sensitive to metabolic alteration *in vivo*. **A.** Body mass, HbA1c, GAL3, and NOX1 mRNA levels in freshly isolated mesenteric endothelial cells from male 20-week-old H*_db_*W*_Nox1_*, H*_db_*K*_Nox1,_* K*_db_*W*_Nox1_,* and K*_db_*K*_Nox1_* (n=5-9). **B.** Body mass, HbA1c, GAL3, and NOX1 mRNA levels in freshly isolated mesenteric endothelial cells from male 20-week-old H*_db_*H*_Mstn_*, H*_db_*K*_Mstn_*, K*_db_*H*_Mstn_,* and K*_db_*K*_Mstn_* (n=4-6). **C.** Body mass, HbA1c, GAL3, and NOX1 mRNA levels in freshly isolated mesenteric endothelial cells from male 20-week-old H*_db_*H*_Ptp1b_*, H*_db_*K*_Ptp1b_*, K*_db_*H*_Ptp1b_,* and K*_db_*K*_Ptp1b_* (n=4-6). **D.** Body mass, HbA1c, GAL3, and NOX1 mRNA levels in freshly isolated mesenteric endothelial cells from male 20-week-old H*_db_* and K*_db_* mice with or without metformin treatment (n=3). All data are represented as mean ± SEM.

## Discussion

Endothelial dysfunction is a classical early mechanism in the development of vascular disease like hypertension, likely driven by a pro-oxidant and pro-inflammatory insult to the endothelium.^2,3^ Metabolic dysfunction is well-known to contribute to endothelial dysfunction, but identification of a mechanism linking aberrant metabolism and cardiovascular function has proven elusive. In the present study, we present evidence that the sugar binding lectin, GAL3, is overexpressed in the microvascular endothelium in obesity. We demonstrate that deletion of GAL3 causes no significant alterations in overall phenotype or metabolism in lean and obese mice, except for resolving plasma ROS production to levels seen with lean controls. GAL3 deletion rescued endothelial-dependent vasodilation and hypertension, most likely by decreasing the expression of NOX1-derived ROS in parallel with resolved inflammation. Finally, we demonstrate that GAL3 expression is sensitive to improvement in glucose metabolism through treatment with metformin, improvement of insulin signaling, or augmentation of muscle mass.

GAL3 is a carbohydrate binding lectin that has been demonstrated to play a significant role mechanisms governing inflammation, proliferation, migration, and apoptosis.^24–26,42–43^ Overexpression of GAL3 has been well-demonstrated in obesity, as well as in many other inflammatory diseases, but is best studied in cancer and pulmonary hypertension.^25,44–46^ Cardiac GAL3 is also an early biomarker for heart failure, and contributes to the development of cardiac inflammation and fibrosis.^47–48^ While the role of GAL3 in promoting inflammation and proliferation in vascular smooth muscle has been further explored in recent studies, the mechanistic contributions of endothelial GAL3 to vascular disease remain largely unexplored. In this study, we report the first evidence that endothelial GAL3 directly contributes to the pathology of microvascular dysfunction in obesity, and that deletion of GAL3 rescues both endothelium-dependent vasodilation and hypertension in obese mice to that of lean controls. The role of GAL3 was exclusively studied in male mice since female *db/db* mice are not overtly hyperglycemic and do not develop endothelial dysfunction to the degree of male *db/db* mice. We would hypothesize that since female mice are normoglycemic and GAL3 is glucose-sensitive, GAL3 is not upregulated and does not mediate endothelial function in female *db/db* mice. Human data were divided by sex and analyzed, however no significant differences in plasma GAL3 levels were detected. These data establish GAL3 as a mechanistic driver of vascular dysfunction in metabolic disease and an attractive target for future therapeutic modulation in obese individuals.

We and others have consistently demonstrated that NOX1 is the major ROS-contributing enzyme contributing to pathology in the vasculature of obese mice, as other NOX isoforms are essential in maintaining proper signaling in the endothelium and smooth muscle.^7^ For example, NOX4 has been shown to mediate both beneficial and detrimental effects, perhaps making it a difficult target for therapeutic intervention.^49^ NOX2 expression is largely limited to macrophages, supporting the role of NOX1 as the most prominent pathological isoform in the vascular endothelium.^50^ In this study, we demonstrate that GAL3 regulates endothelial NOX1 expression in obesity, without altering expression of these other NOX isoforms. Promoter luciferase assays suggest that GAL3 can increase NOX1 promoter activity and while the mechanisms involved are not yet defined, they will be an important focus for future studies. Taken together, these data indicate a novel, mechanistic role for GAL3 in increasing endothelial ROS production, most likely by driving the overexpression of NOX1. While the deleterious effects of ROS overproduction are well-understood, therapeutics such as antioxidants seeking to abrogate ROS production and accumulation have been largely unsuccessful, necessitating the identification of an upstream target that regulates superoxide-generating enzymes to produce the desired effect and preserve vascular health^15^. We propose that GAL3 is such a target, and that pharmacological inhibition of GAL3 is deserving of future study.

Our data indicate that GAL3 expression is driven by to beneficial alterations in metabolism, specifically by improving glucose handling. Previous work from our laboratory has demonstrated that increasing muscle mass in obesity significantly improves both cardiovascular and renal function, highlighting the importance of preserving healthy skeletal muscle.^8,40–41^ Indeed, muscle mass is negatively correlated with cardiovascular outcomes in humans, as sarcopenia associated with aging and sedentary lifestyle is an independent predictor of cardiovascular morbidity and mortality.^51–54^ Published data indicate that in obese animals lacking myostatin, a potent negative regulator of muscle growth, endothelial function is improved by restoring healthy eNOS/NOX1 balance.^8,40–41^ Preserved skeletal muscle ameliorates obesity-induced hyperglycemia, thereby reducing the amount of glycated end-products that would otherwise elicit pathological downstream signaling by GAL3.^8^ Further supporting this key finding are our results that improvement of glucose metabolism by deletion of PTP1B or treatment with metformin causes a corresponding decrease in endothelial GAL3/NOX1 expression. Whether metformin improves GAL3 signaling by direct effects on the endothelium or by the indirect effect of lowering blood glucose remains to be determined. However, since the drug is so well-tolerated and widely available, these data provide compelling rationale that low-dose treatment with metformin in obesity, even before the onset of diabetes or cardiovascular complications, may prove beneficial in ameliorating pathological expression of GAL3 and subsequent entry into cardiometabolic disease.

In summary, we demonstrate that endothelial GAL3 mediates obesity-induced microvascular dysfunction and hypertension and regulates NOX1 expression and subsequent overproduction of ROS. Inability to properly maintain microvascular function is an early hallmark of cardiovascular disease and accelerates end-organ damage due to loss of proper tissue perfusion. However, GAL3 expression is sensitive to improvement in glucose handling by treatment with metformin, improvement of insulin signaling, and augmenting muscle mass, indicating that GAL3 is not only a biomarker of vascular disease, but rather plays a mechanistic role in its development by acting as a metabolic sensor between aberrant metabolism and vascular dysfunction. These data indicate that inhibition of GAL3 is a novel target for therapeutic intervention to ameliorate cardiovascular disease in obese individuals, thereby improving indices of morbidity and mortality.

## Abbreviations

Ach: Acetylcholine
AGER: Receptor for advanced glycation end-products
AgRP: Agouti-related peptide
BGAL: Beta-galactosidase
eNOS: Endothelial nitric oxide synthase
GAL3: Galectin-3
HEK cells: Human embryonic kidney cells
HSP90: Heat shock protein 90
KCl: Potassium chloride
KO: Knockout
MAP: Mean arterial pressure
Met: Metformin
NO: Nitric oxide
NOX: Nicotinamide adenine dinucleotide phosphate oxidase
OLR1: Oxidized low density lipoprotein receptor 1
PE: Phenylephrine
PTP1B: Protein tyrosine phosphatase 1b
RAGE: Receptor for advanced glycation end-products
ROS: Reactive oxygen species
SNP: Sodium nitroprusside
T2DM: Type II Diabetes Mellitus
TBARS: Thiobarbituric acid reactive substances

## Acknowledgments

We would like to thank Kristen Carver and the Augusta University Electron Microscopy and Histology Core for their assistance with specimen preparation, sectioning, and mounting. We also thank Safia Ogbi for assistance with Western Blotting. Visual aids were created with BioRender.com. Finally, we are grateful to the reviewers for taking time to provide thought-provoking criticism and insightful suggestions to improve the rigor and reproducibility of this study.

## References

1. van Rooy MJ, Pretorius E. Obesity, hypertensin and hypercholesterolemia as risk factors for atherosclerosis leading to ischemic events. Curr Med Chem. 2014;21(19):2121–9. doi:10.2174/0929867321666131227162950

2. Konukoglu D, Uzun H. Endothelial Dysfunction and Hypertension. Adv Exp Med Biol. 2017;956:511–540. doi:10.1007/5584_2016_90

3. Suzuki T, Hirata K, Elkind MS, et al. Metabolic syndrome, endothelial dysfunction, and risk of cardiovascular events: the Northern Manhattan Study (NOMAS). Am Heart J. 2008;156(2):405–410. doi:10.1016/j.ahj.2008.02.022

4. Cyr AR, Huckaby LV, Shiva SS, Zuckerbraun BS. Nitric Oxide and Endothelial Dysfunction. Crit Care Clin. 2020;36(2):307–321. doi:10.1016/j.ccc.2019.12.009

5. Incalza MA, D’Oria R, Natalicchio A, Perrini S, Laviola L, Giorgino F. Oxidative stress and reactive oxygen species in endothelial dysfunction associated with cardiovascular and metabolic diseases. Vascul Pharmacol. 2018;100:1–19. doi:10.1016/j.vph.2017.05.005

6. Förstermann U, Xia N, Li H. Roles of Vascular Oxidative Stress and Nitric Oxide in the Pathogenesis of Atherosclerosis. Circ Res. 2017;120(4):713–735. doi:10.1161/CIRCRESAHA.116.309326

7. Thompson JA, Larion S, Mintz JD, Belin de Chantemèle EJ, Fulton DJ, Stepp DW. Genetic Deletion of NADPH Oxidase 1 Rescues Microvascular Function in Mice With Metabolic Disease. Circ Res. 2017;121(5):502–511. doi:10.1161/CIRCRESAHA.116.309965

8. Qiu S, Mintz JD, Salet CD, et al. Increasing muscle mass improves vascular function in obese (db/db) mice. J Am Heart Assoc. 2014;3(3):e000854. Published 2014 Jun 25. doi:10.1161/JAHA.114.000854

9. Muñoz M, López-Oliva ME, Rodríguez C, et al. Differential contribution of Nox1, Nox2 and Nox4 to kidney vascular oxidative stress and endothelial dysfunction in obesity. Redox Biol. 2020;28:101330. doi:10.1016/j.redox.2019.101330

10. Gimenez M, Schickling BM, Lopes LR, Miller FJ Jr. Nox1 in cardiovascular diseases: regulation and pathophysiology. Clin Sci (Lond). 2016;130(3):151–165. doi:10.1042/CS20150404

11. San Martín A, Du P, Dikalova A, et al. Reactive oxygen species-selective regulation of aortic inflammatory gene expression in Type 2 diabetes. Am J Physiol Heart Circ Physiol. 2007;292(5):H2073–H2082. doi:10.1152/ajpheart.00943.2006

12. Santhanam AV, d’Uscio LV, Smith LA, Katusic ZS. Uncoupling of eNOS causes superoxide anion production and impairs NO signaling in the cerebral microvessels of hph-1 mice. J Neurochem. 2012;122(6):1211–1218. doi:10.1111/j.1471-4159.2012.07872.x

13. Yang YM, Huang A, Kaley G, Sun D. eNOS uncoupling and endothelial dysfunction in aged vessels. Am J Physiol Heart Circ Physiol. 2009;297(5):H1829–H1836. doi:10.1152/ajpheart.00230.2009

14. Meza CA, La Favor JD, Kim DH, Hickner RC. Endothelial Dysfunction: Is There a Hyperglycemia-Induced Imbalance of NOX and NOS?. Int J Mol Sci. 2019;20(15):3775. Published 2019 Aug 2. doi:10.3390/ijms20153775

15. Radi R. Oxygen radicals, nitric oxide, and peroxynitrite: Redox pathways in molecular medicine. Proc Natl Acad Sci U S A. 2018;115(23):5839–5848. doi:10.1073/pnas.1804932115

16. Ye Y, Li J, Yuan Z. Effect of antioxidant vitamin supplementation on cardiovascular outcomes: a meta-analysis of randomized controlled trials. PLoS One. 2013;8(2):e56803. doi:10.1371/journal.pone.0056803

17. Heart Outcomes Prevention Evaluation Study Investigators, Yusuf S, Dagenais G, Pogue J, Bosch J, Sleight P. Vitamin E supplementation and cardiovascular events in high-risk patients. N Engl J Med. 2000;342(3):154–160. doi:10.1056/NEJM200001203420302

18. Fox CS, Coady S, Sorlie PD, et al. Increasing cardiovascular disease burden due to diabetes mellitus: the Framingham Heart Study. Circulation. 2007;115(12):1544–1550. doi:10.1161/CIRCULATIONAHA.106.658948

19. Schmidt AM. Diabetes Mellitus and Cardiovascular Disease. Arterioscler Thromb Vasc Biol. 2019;39(4):558–568. doi:10.1161/ATVBAHA.119.310961

20. Egaña-Gorroño L, López-Díez R, Yepuri G, et al. Receptor for Advanced Glycation End Products (RAGE) and Mechanisms and Therapeutic Opportunities in Diabetes and Cardiovascular Disease: Insights From Human Subjects and Animal Models. Front Cardiovasc Med. 2020;7:37. Published 2020 Mar 10. doi:10.3389/fcvm.2020.00037

21. Nogueira-Machado JA, Chaves MM. From hyperglycemia to AGE-RAGE interaction on the cell surface: a dangerous metabolic route for diabetic patients. Expert Opin Ther Targets. 2008;12(7):871–882. doi:10.1517/14728222.12.7.871

22. Soro-Paavonen A, Watson AM, Li J, et al. Receptor for advanced glycation end products (RAGE) deficiency attenuates the development of atherosclerosis in diabetes. Diabetes. 2008;57(9):2461–2469. doi:10.2337/db07-1808

23. Reiniger N, Lau K, McCalla D, et al. Deletion of the receptor for advanced glycation end products reduces glomerulosclerosis and preserves renal function in the diabetic OVE26 mouse. Diabetes. 2010;59(8):2043–2054. doi:10.2337/db09-1766

24. Hudson BI, Lippman ME. Targeting RAGE Signaling in Inflammatory Disease. Annu Rev Med. 2018;69:349–364. doi:10.1146/annurev-med-041316-085215

25. Sciacchitano S, Lavra L, Morgante A, et al. Galectin-3: One Molecule for an Alphabet of Diseases, from A to Z. Int J Mol Sci. 2018;19(2):379. Published 2018 Jan 26. doi:10.3390/ijms19020379

26. Menini S, Iacobini C, Blasetti Fantauzzi C, Pesce CM, Pugliese G. Role of Galectin-3 in Obesity and Impaired Glucose Homeostasis. Oxid Med Cell Longev. 2016;2016:9618092. doi:10.1155/2016/9618092

27. Pugliese G, Iacobini C, Pesce CM, Menini S. Galectin-3: an emerging all-out player in metabolic disorders and their complications. Glycobiology. 2015;25(2):136–150. doi:10.1093/glycob/cwu111

28. Gao Z, Liu Z, Wang R, Zheng Y, Li H, Yang L. Galectin-3 Is a Potential Mediator for Atherosclerosis. J Immunol Res. 2020;2020:5284728. Published 2020 Feb 14. doi:10.1155/2020/5284728

29. Oikonomou E, Zografos T, Papamikroulis GA, et al. Biomarkers in Atrial Fibrillation and Heart Failure. Curr Med Chem. 2019;26(5):873–887. doi:10.2174/0929867324666170830100424

30. Amin HZ, Amin LZ, Wijaya IP. Galectin-3: a novel biomarker for the prognosis of heart failure. Clujul Med. 2017;90(2):129–132. doi:10.15386/cjmed-751

31. Challis RC, Ravindra Kumar S, Chan KY, et al. Systemic AAV vectors for widespread and targeted gene delivery in rodents [published correction appears in Nat Protoc. 2019 Aug;14(8):2597]. Nat Protoc. 2019;14(2):379–414. doi:10.1038/s41596-018-0097-3

32. Placeholder for accompanying manuscript Sellers & Padgett et al.

33. Valente AJ, Zhou Q, Lu Z, et al. Regulation of NOX1 expression by GATA, HNF-1alpha, and Cdx transcription factors. Free Radic Biol Med. 2008;44(3):430–443. doi:10.1016/j.freeradbiomed.2007.10.035

34. Chen F, Li X, Aquadro E, et al. Inhibition of histone deacetylase reduces transcription of NADPH oxidases and ROS production and ameliorates pulmonary arterial hypertension. Free Radic Biol Med. 2016;99:167–178. doi:10.1016/j.freeradbiomed.2016.08.003

35. Zhao Z, Xu X, Cheng H, et al. Galectin-3 N-terminal tail prolines modulate cell activity and glycan-mediated oligomerization/phase separation. Proc Natl Acad Sci U S A. 2021;118(19):e2021074118. doi:10.1073/pnas.2021074118

36. Diamant M, Tushuizen ME. The metabolic syndrome and endothelial dysfunction: common highway to type 2 diabetes and CVD [published correction appears in Curr Diab Rep. 2006 Dec;6(6):iii]. Curr Diab Rep. 2006;6(4):279–286. doi:10.1007/s11892-006-0061-4

37. Caballero AE, Arora S, Saouaf R, et al. Microvascular and macrovascular reactivity is reduced in subjects at risk for type 2 diabetes. Diabetes. 1999;48(9):1856–1862. doi:10.2337/diabetes.48.9.1856

38. Zhang Y, Hu M, Ma H, et al. The impairment of reproduction in db/db mice is not mediated by intraovarian defective leptin signaling. Fertil Steril. 2012;97(5):1183–1191. doi:10.1016/j.fertnstert.2012.01.126

39. Ali MI, Ketsawatsomkron P, Belin de Chantemele EJ, et al. Deletion of protein tyrosine phosphatase 1b improves peripheral insulin resistance and vascular function in obese, leptin-resistant mice via reduced oxidant tone. Circ Res. 2009;105(10):1013–1022. doi:10.1161/CIRCRESAHA.109.206318

40. Butcher JT, Mintz JD, Larion S, et al. Increased Muscle Mass Protects Against Hypertension and Renal Injury in Obesity. J Am Heart Assoc. 2018;7(16):e009358. doi:10.1161/JAHA.118.009358

41. Butcher JT, Ali MI, Ma MW, et al. Effect of myostatin deletion on cardiac and microvascular function. Physiol Rep. 2017;5(23):e13525. doi:10.14814/phy2.13525

42. Henderson NC, Sethi T. The regulation of inflammation by galectin-3. Immunol Rev. 2009;230(1):160–171. doi:10.1111/j.1600-065X.2009.00794.x

43. Dumic J, Dabelic S, Flögel M. Galectin-3: an open-ended story. Biochim Biophys Acta. 2006;1760(4):616–635. doi:10.1016/j.bbagen.2005.12.020

44. Song L, Tang JW, Owusu L, Sun MZ, Wu J, Zhang J. Galectin-3 in cancer. Clin Chim Acta. 2014;431:185–191. doi:10.1016/j.cca.2014.01.019

45. Barman SA, Li X, Haigh S, et al. Galectin-3 is expressed in vascular smooth muscle cells and promotes pulmonary hypertension through changes in proliferation, apoptosis, and fibrosis. Am J Physiol Lung Cell Mol Physiol. 2019;316(5):L784–L797. doi:10.1152/ajplung.00186.2018

46. Fulton DJR, Li X, Bordan Z, et al. Galectin-3: A Harbinger of Reactive Oxygen Species, Fibrosis, and Inflammation in Pulmonary Arterial Hypertension. Antioxid Redox Signal. 2019;31(14):1053–1069. doi:10.1089/ars.2019.7753

47. Martínez-Martínez E, Brugnolaro C, Ibarrola J, et al. CT-1 (Cardiotrophin-1)-Gal-3 (Galectin-3) Axis in Cardiac Fibrosis and Inflammation. Hypertension. 2019;73(3):602–611. doi:10.1161/HYPERTENSIONAHA.118.11874

48. Dong R, Zhang M, Hu Q, et al. Galectin-3 as a novel biomarker for disease diagnosis and a target for therapy (Review). Int J Mol Med. 2018;41(2):599–614. doi:10.3892/ijmm.2017.3311

49. Schröder K, Zhang M, Benkhoff S, et al. Nox4 is a protective reactive oxygen species generating vascular NADPH oxidase. Circ Res. 2012;110(9):1217–1225. doi:10.1161/CIRCRESAHA.112.267054

50. Singel KL, Segal BH. NOX2-dependent regulation of inflammation. Clin Sci (Lond). 2016;130(7):479–490. doi:10.1042/CS20150660

51. Warren TY, Barry V, Hooker SP, Sui X, Church TS, Blair SN. Sedentary behaviors increase risk of cardiovascular disease mortality in men. Med Sci Sports Exerc. 2010;42(5):879–885. doi:10.1249/MSS.0b013e3181c3aa7e

52. Chin SO, Rhee SY, Chon S, et al. Sarcopenia is independently associated with cardiovascular disease in older Korean adults: the Korea National Health and Nutrition Examination Survey (KNHANES) from 2009. PLoS One. 2013;8(3):e60119. doi:10.1371/journal.pone.0060119

53. Xia MF, Chen LY, Wu L, et al. Sarcopenia, sarcopenic overweight/obesity and risk of cardiovascular disease and cardiac arrhythmia: A cross-sectional study. Clin Nutr. 2021;40(2):571–580. doi:10.1016/j.clnu.2020.06.003

54. Lavie CJ, Ozemek C, Carbone S, Katzmarzyk PT, Blair SN. Sedentary Behavior, Exercise, and Cardiovascular Health. Circ Res. 2019;124(5):799–815. doi:10.1161/CIRCRESAHA.118.312669

